# Mutation-specific reporter for the optimization and enrichment of prime editing

**DOI:** 10.1101/2021.05.08.443062

**Authors:** I.F. Schene, I.P. Joore, J.H.L. Baijens, S. Shehata, E.F. Ilcken, D.P. Bolhuis, R.C.M. van Rees, S.A. Spelier, P.J. van der Doef, J.M. Beekman, E.E.S. Nieuwenhuis, S.A. Fuchs

## Abstract

We present a fluorescent prime editing and enrichment reporter (fluoPEER), which can be tailored to any genomic target site. This system rapidly and faithfully ranks the efficiency of prime edit guide RNAs (pegRNAs) and any prime editor protein, including novel variants with flexible PAM recognition. Successful reporter editing enriches for genomic editing. FluoPEER can be employed for efficient correction of patient cells and to elucidate cellular mechanisms needed for successful prime editing.

## Main

Prime editing is a precise genome-editing technique to generate any moderately sized mutation without the need for double strand breaks (Anzalone, 2019). This technique enables repair of patient mutations, the generation of disease models, and in vivo editing (Schene, 2020; Liu, 2021). The requisite NGG-PAM sequence near edit sites limits the capacity of the original prime editor (PE2) to generate mutations. PE2 variants with flexible PAM recognition (Nishimasu, 2018; Walton, 2018; Kweon, 2021) provide sufficient PAM-sites to target any smaller pathogenic mutation included in the ClinVar database (Supplementary Fig. 1).

The mechanisms required for efficient prime editing are poorly understood. More specifically, optimal design of the prime edit guide RNA (pegRNA) remains largely elusive. However, it is known that primer binding site (PBS) and reverse transcriptase template (RTT) variations have marked effects on editing efficiency (Anzalone, 2019; Lin, 2021). A deep learning strategy has further specified important pegRNA characteristics, which can now be used to instruct ‘NGG’ pegRNA design (Kim, 2021). However, this deep learning algorithm failed to predict a pegRNA efficiency score for >80% of the mutations in the ClinVar database (Supplementary Fig. 2b). As expected, for those pathogenic variants within the scope of the prediction algorithm, a higher number of available PAMs per mutation resulted in a higher predicted maximum pegRNA efficiency (Supplementary Fig. 2c). To address the shortcomings of current prediction methods and harness the strength of new PE2 variants, we present a fluorescent prime editing and enrichment reporter, called fluoPEER. This reporter can be tailored to any genomic target site and allows high-throughput analysis using fluorescence-activated cell sorting (FACS) within seven days from starting cloning.

We cloned genomic target regions (54-70 nucleotides) spanning all possible pegRNAs between an eGFP and a Cherry cassette under constitutive expression (Fig. 1a). When the target mutation did not contain a native stop-codon or frameshift, a stop-codon or frameshift was inserted (Fig. 1a). We hypothesized that this adaptation does not limit predictive power since the substitution or insertion of single nucleotides in the RTT-region has minor effects on prime editing efficiency (Supplementary Fig. 3). Importantly, this system allows targeting of the reporter and the corresponding genomic locus with the same pegRNA design.

**Figure 1:**
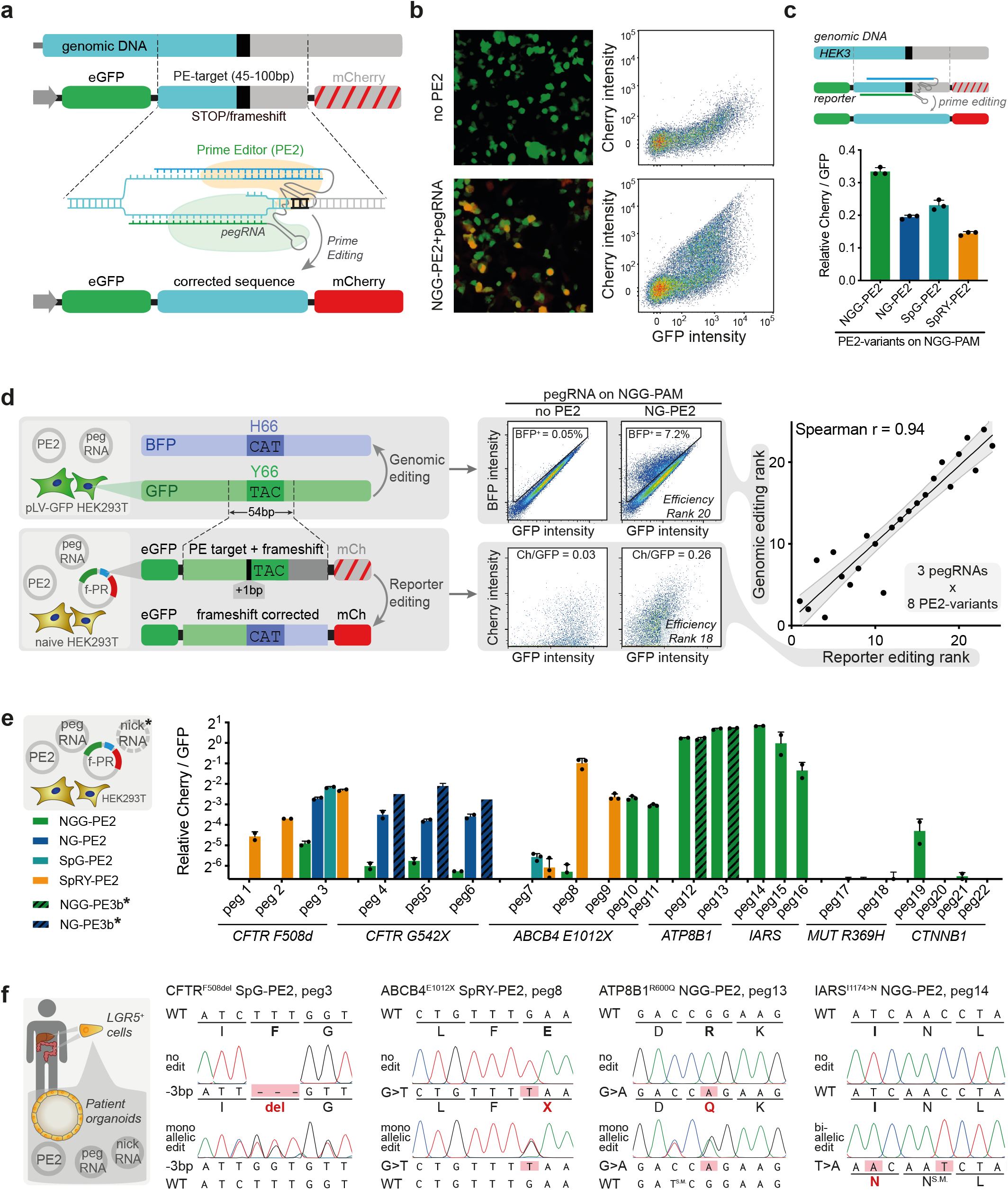
A fluorescence-based prime-editing reporter system instructs pegRNA design for genomic editing. **a**, The fluoPEER plasmid uses a 50-70 bp genomic region containing a stop codon or frameshift. In case the genomic region does not contain a naturally occurring stop codon or frameshift, one is added. The prime edit machinery targets and edits the genomic insert, removing the insertion or stop codon. The same prime edit machinery, including the same pegRNA design, is capable of editing the genomic DNA. A finished fluoPEER plasmid contains a GFP-P2A-Genomic insert-P2A-Cherry. **b**, Editing of the genomic insert is visualized by Cherry signal and quantified using flow cytometry. **c**, fluoPEER distinguishes between the efficiency of different PE2 variants. This is quantified as the total Cherry signal over the total GFP signal, which gives a measure of editing per transfected plasmid. **d**, A comparison between the generation of a GFP to BFP conversion using prime editing on the genomic DNA and on the fluoPEER plasmid. HEK293T cells containing a lentivirally integrated genomic GFP cassette are transfected with the prime editing machinery to convert GFP to BFP in 24 conditions total, while the same conditions are applied to HEK293T cells cotransfected with the fluoPEER plasmid for the GFP to BFP conversion. Comparing the efficiency ranking of editing conditions, a strong spearman correlation is found (0.94) between fluoPEER and genomic DNA editing. **e**, Prime editing on fluoPEER yields efficiency read-outs for 7 edits for 22 pegRNAs in total. **f**, Using the optimal pegRNAs based on fluoPEER data, several pathogenic mutations in patient-derived organoids are genetically corrected and organoids with biallelic *IARS1* mutations are generated in wildtype liver organoids.

We transfected fluoPEER together with prime editing machinery in HEK293T cells and analyzed reporter editing after 3 days using FACS (Fig. 1b). Based on the ratio of Cherry to GFP signal, an efficiency score was established for each editing condition, enabling ranking of PE2 variants and pegRNAs (Fig. 1c). To verify the reliability of this reporter system, we used the same combinations of PE2 variants and pegRNAs to convert a genomically integrated GFP gene to BFP. When ranking the editing conditions based on efficiency for both strategies, correlation between genomic editing and reporter prediction was very strong (r=0.94, Fig. 1d, Supplementary Fig. 4). This high correlation further confirms that the additional insertion of a single nucleotide in the reporter does not hamper predictive capacity. Importantly, Sanger sequencing could only detect genomic editing in conditions with >10% BFP+ cells (Supplementary Fig. 4), supporting the need for a sensitive reporter system when targeting difficult-to-edit loci.

Next, we compared the DeepPE algorithm to fluoPEER. In contrast to the algorithm, fluoPEER ranked pegRNA efficiencies correctly at each genomic locus where the deep learning tool could not (Supplementary Fig. 5). Furthermore, we confirmed the robustness of the Cherry over GFP ratio as a read-out for efficiency by varying the input reporter plasmid concentration (Supplementary Fig. 6). By varying the time point of read-out, we further corroborated stability of the ratio-based ranking was stable over a period of 6 days (Supplementary Fig. 6).

Using the reporter system, we tested pegRNAs and PE2 variant combinations for many different genomic loci within 7 days from the start of cloning (Fig. 1e). Selecting the strategy with the highest reporter prediction score enabled quick and efficient repair of disease-causing mutations in patient-derived organoids (Fig. 1f). These included common mutations causing cystic fibrosis (*CFTR*^F508del^ and *CFTR*^G542X^), progressive familial intrahepatic cholestasis (PFIC) type 3 (*ABCB4*^E1012X^), and PFIC type 1 (*ATP8B1*^R600Q^). Furthermore, we used fluoPEER ranking to efficiently mutate cytosolic isoleucyl-tRNA synthetase (*IARS1*) in liver-derived organoids. This confirmed that biallelic, but not monoallelic, *IARS1*^I1174N^ mutations impaired organoid expandability, reflecting the failure to thrive observed in patients with biallelic *IARS1* mutations (Kopajtich, 2016; Fuchs, 2019; Supplementary Fig. 7). Importantly, pegRNAs targeting a difficult-to-edit locus to repair a common methylmalonic acidemia mutation (*MUT*^R369H^) did not edit on the reporter and also failed in organoids, demonstrating accurate negative prediction. Lastly, fluoPEER correctly predicted pegRNA efficiencies to edit *CTNNB1* as observed in earlier organoid experiments (Schene, 2020) (Supplementary Fig 7).

To further characterize new PE2 variants, we adapted the genomic target region of fluoPEER to test the PAM-specificities of each flexible prime editor. We found that the SpG- and NG-PE2 complexes have very similar flexible PAM preferences with the SpG showing higher efficiency scores overall (Supplementary Fig. 8a-b). The SpRY-PE2 displayed a PAM-specificity pattern highly similar to the SpRY-Cas9 protein (Walton et al., 2020), essentially functioning as a PAM-less prime editor. Interestingly, a guanine nucleotide on the 4th PAM position led to higher editing efficiency compared to cytosine (Supplementary Fig. 8b). We further tested PE*-variants with improved nuclear localization (Liu, 2021), but this did not improve efficiency of reporter editing and genomic editing compared to standard PE2 variants (Supplementary Fig. 8c-d).

Selection of cells with a detectable edit has been shown to enrich for a simultaneously introduced edit at another locus (Li, 2021). To test whether fluoPEER allows such enrichment, we sorted transfected cells that were plasmid-edited (GFP^+^Cherry^+^) and found that these cells showed large increases (>2 fold) in genomic editing compared to plasmid-unedited cells (GFP^+^Cherry^-^) (Fig. 2a). The same was observed for editing in patient-derived organoids, corroborating that this system can function as an enrichment tool for the repair or generation of patient mutations (Fig. 2a).

**Figure 2:**
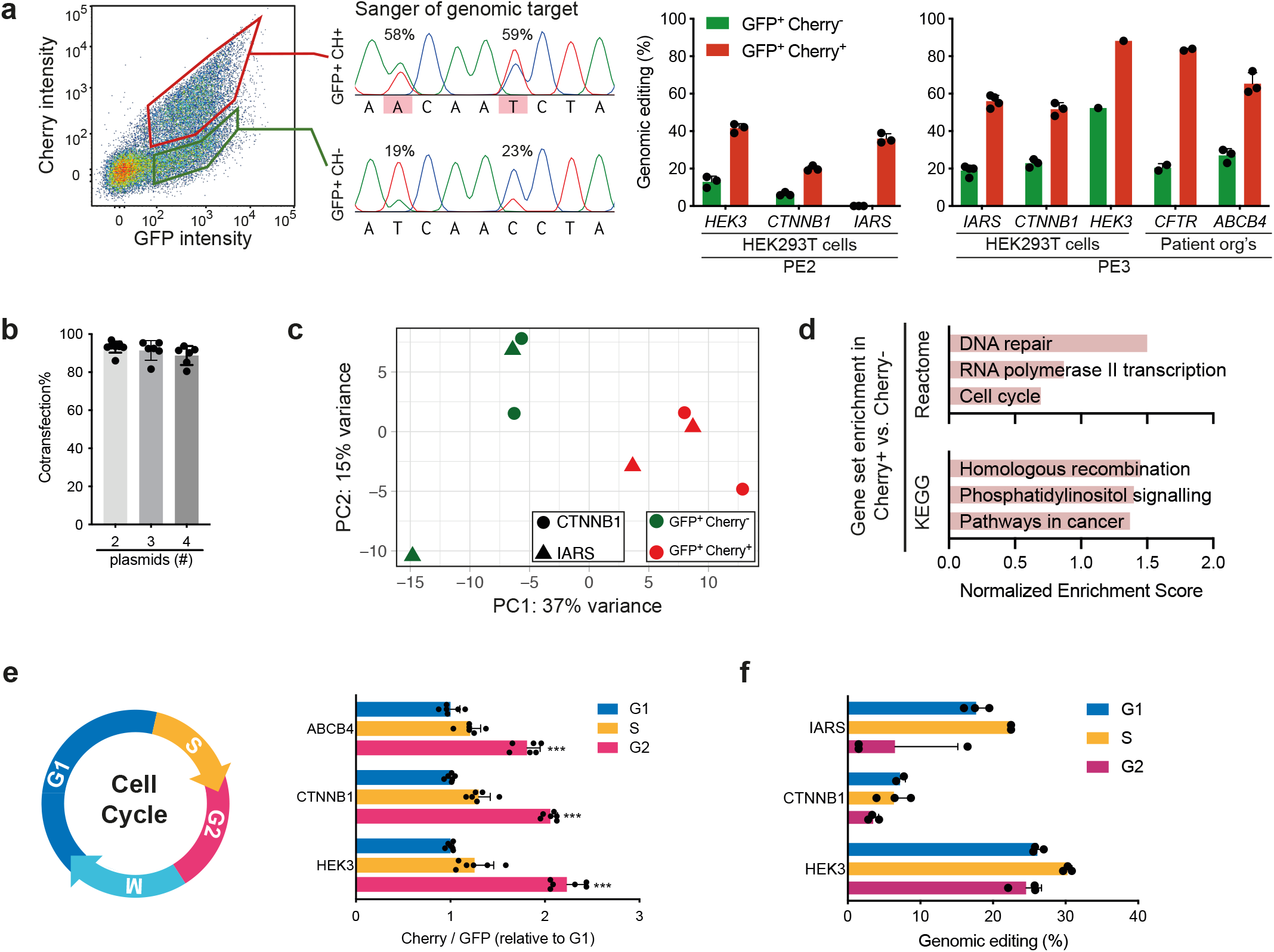
fluoPEER enriches for genomic editing. **a**, Sorting based on presence (GFP^+^Cherry^+^) and absence (GFP^+^Cherry^-^) of reporter editing shows a strong enrichment of genomic editing in the reporter edited population in HEK293T cells and organoids. Cells were sorted 24 (PE2) or 72 (PE3) hours after transfection. **b**, Cotransfection of 2, 3, or 4 different fluorescent plasmids shows that >90% of the cells successfully transfected with a single plasmid also received all other plasmids. **c**, PCA plot and **d**, gene set enrichment analysis of RNA sequencing of the GFP^+^Cherry^+^ and GFP^+^Cherry^-^ populations show expression-based differences in DNA repair- and cell cycle-associated genes. **e**, Hoechst staining in HEK293T cells transfected with the fluoPEER plasmid and prime editing machinery shows a strong enrichment of reporter editing signal in G2. **f**, Sanger sequencing of HEK293T cells sorted based on cell cycle shows differences in genomic editing efficiency.

We hypothesized that enrichment of genomic editing was caused by higher transfection rates of the prime editing plasmids in GFP^+^Cherry^+^ cells, but found that >90% of transfected cells received all plasmids (Fig. 2b, Supplementary Fig. 9). This suggested that a subpopulation of cells is more susceptible to prime editing. We therefore compared the transcriptome of reporter-edited versus non-edited cells and found enrichment of DNA repair genes and cell cycle-associated genes (Fig. 2c and 2d, Supplementary Fig. 10). As such, we evaluated the reporter editing of cells during different cell cycle stages using FACS. We found a large increase (>40%) of Cherry signal in cells that were in G2 (Fig. 2e). This indicates an effect of cell cycle-associated mechanisms, but the precise cell cycle stages of importance for prime editing are difficult to infer due to the delay in Cherry expression. We confirmed cell cycle associated differences in prime editing efficiency when looking at genomic DNA editing (Fig. 2f). Due to the time required for gene editing and the concurrent progression in cell cycle, the specific effects of cell cycle stage on prime editing need further elucidation. Nevertheless, these findings indicate a cell cycle dependency for prime editing efficiency. Together, these results establish fluoPEER as a highly dynamic prime editing read-out, enabling enrichment of genomic editing based on plasmid co-editing.

In conclusion, fluoPEER represents a customizable tool for optimization of prime editing strategies and enrichment of edited cells. We demonstrate its strength to optimize pegRNA design and characterize novel PE2 variants for efficient correction of almost any mutation in patient-derived cells. Enrichment of genomic editing by fluoPEER relies on transient selection and does not require transfection of a second pegRNA, genomic integration of a selection cassette, or generation of an additional genomic edit (Katti, 2020; Li, 2021). This versatile tool will facilitate effective prime editing of disease-causing mutations in any relevant cell type.

## Acknowledgements

The authors thank the Kim lab of the Department of Pharmacology, Yonsei University College of Medicine for supplying the PE_prediction code.

## Author contributions

I.F.S., I.P.J. and S.A.F. designed the project; J.H.L.B. analysed the pathogenic mutations in the ClinVar database using PE_prediction; H.P.J.D. helped establishing the biobank of patient-derived stem cell organoids used in this study; I.F.S., I.P.J., S.S., E.F.I., D.P.B. and R.C.M.R performed experiments and analyses; S.A.S. and J.M.B. provided the cystic fibrosis samples used in this study; I.F.S., I.P.J., E.E.S.N., and S.A.F. wrote the manuscript.

## Supplementary figure legends

**Supplementary Figure 1:**
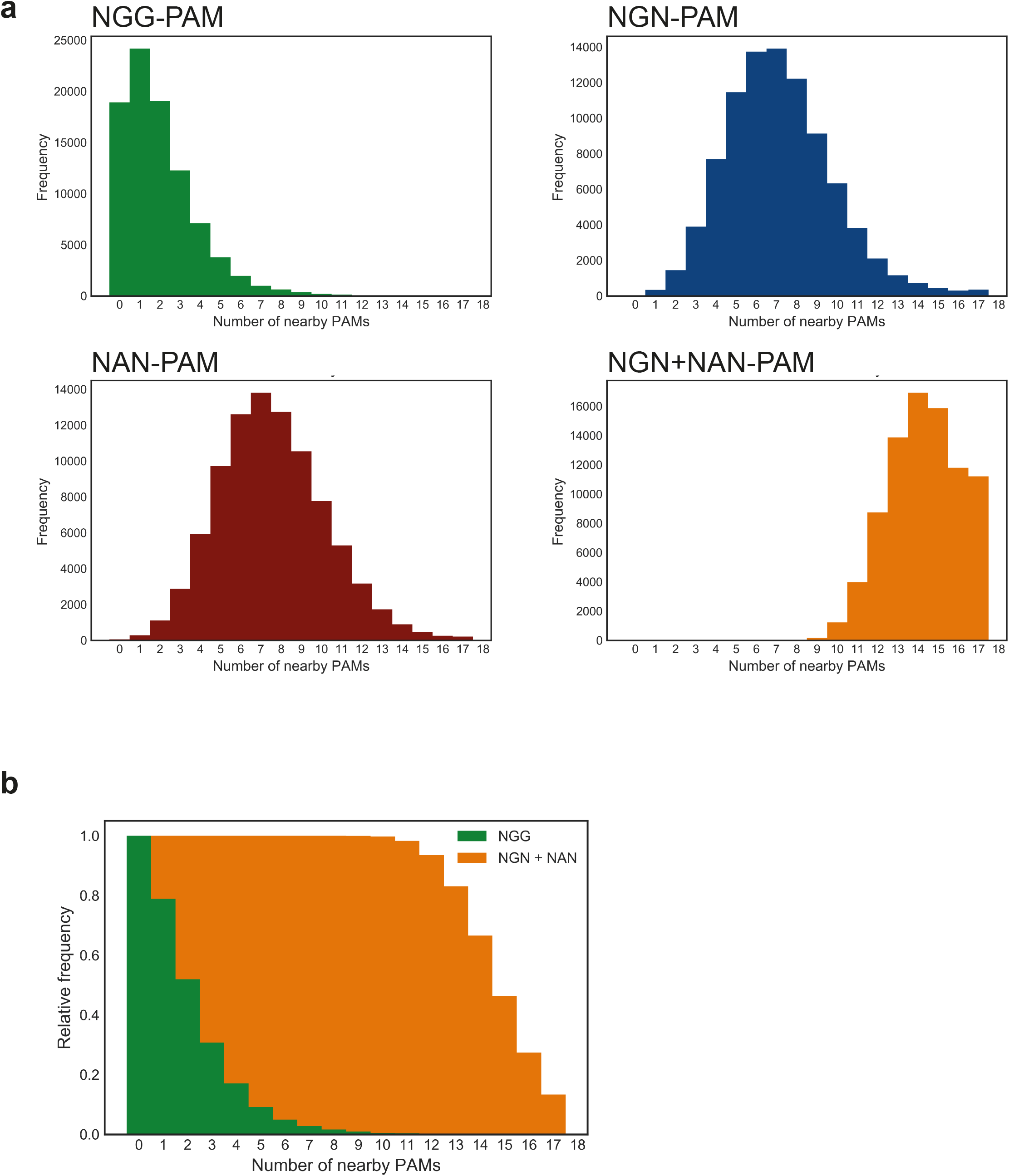
Analysis of the targeting scope of (NGG-)PE2 and flexible PE2 variants within the ClinVar database. **a**, The number of NGG-, NAN-, or NGN-PAMs within the genomic region surrounding pathogenic variants in the ClinVar database. Only mutations shorter than 51 bp were considered as correctable by prime editing and PAM sites in a window of 10bp upstream to 4bp downstream of the target site were considered usable, based on Kim et al., (2021) **b**, Cumulative histogram of the number of available PAMs shows that flexible PE2 variants (NGN+NAN) have at least 9 PAMs available for the repair of ClinVar mutations.

**Supplementary Figure 2:**
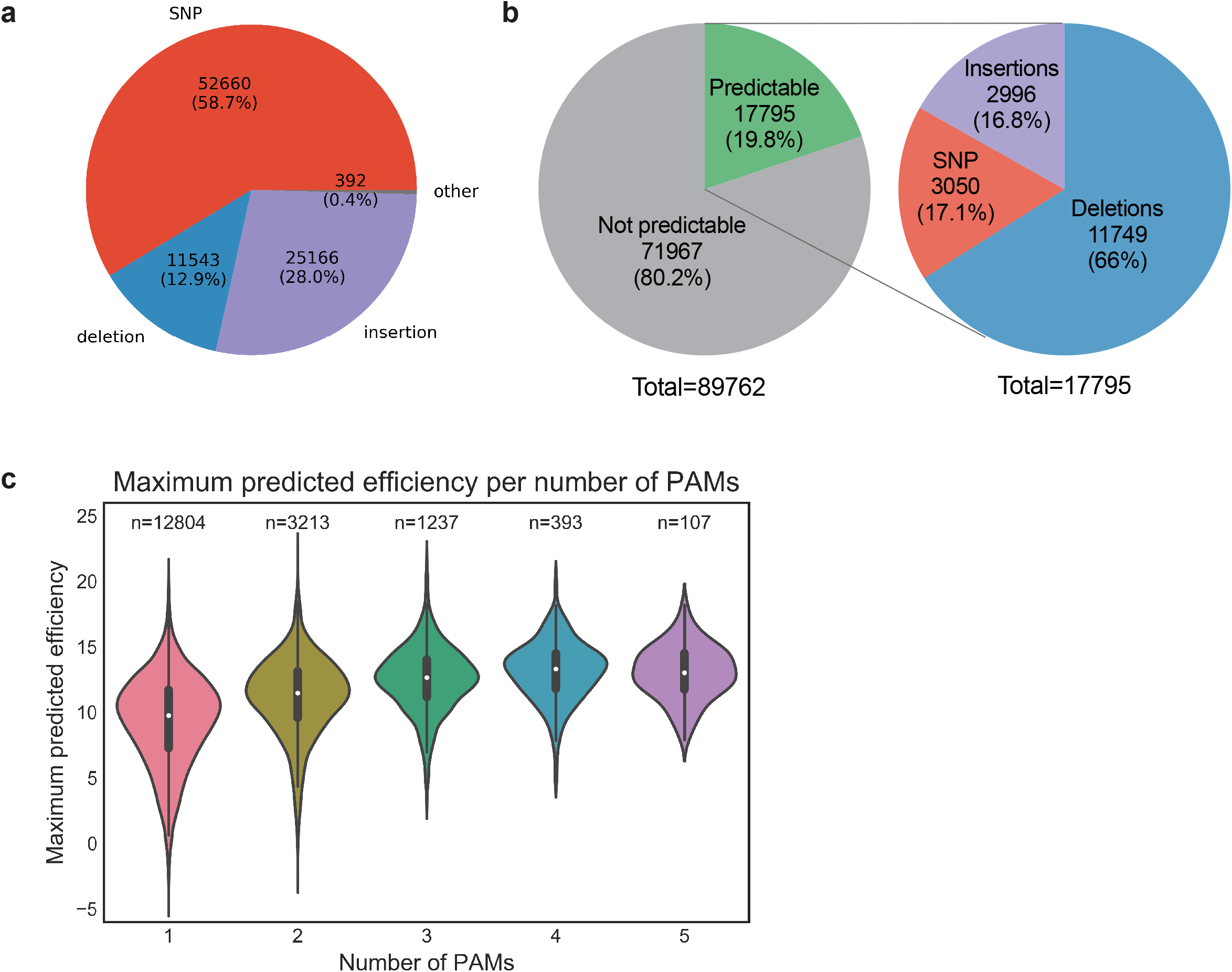
Estimation of reparable mutations in the ClinVar database that are within the scope of the PE_prediction machine learning algorithm. **a**, Types of mutations of pathogenic variants in the ClinVar database **b**, Maximum predicted efficiency for repair of ClinVar pathogenic variants for mutations that fall within the scope of the DeepPE deep learning algorithm. Increase in available PAMs for pathogenic variants results in a higher median maximum predicted repair.

**Supplementary Figure 3:**
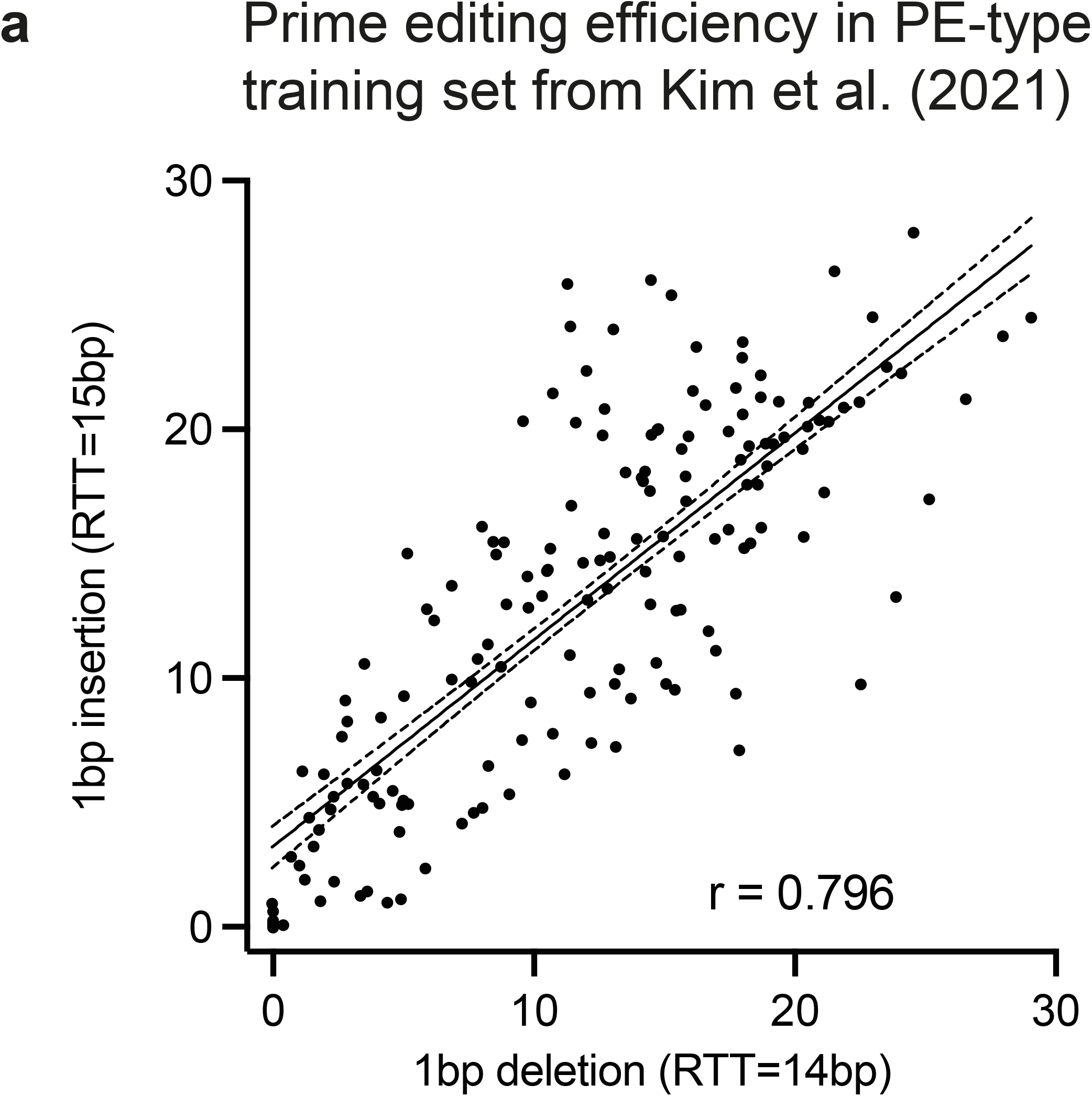
Correlation of prime editing efficiency of 1bp deletions versus 1bp insertions. **a**, Data from Kim et al. (2021) was re-analyzed to estimate the correlation of prime editing efficiencies of pegRNAs that create a 1bp deletion (x-axis) to the efficiencies of corresponding pegRNAs with the same spacer and PBS sequence that create a 1bp insertion. No data was available for the comparison of 1bp deletion- to 1bp substitution-edits since Kim et al. performed 1bp substitutions with much longer RTTs than 1bp deletions.

**Supplementary Figure 4:**
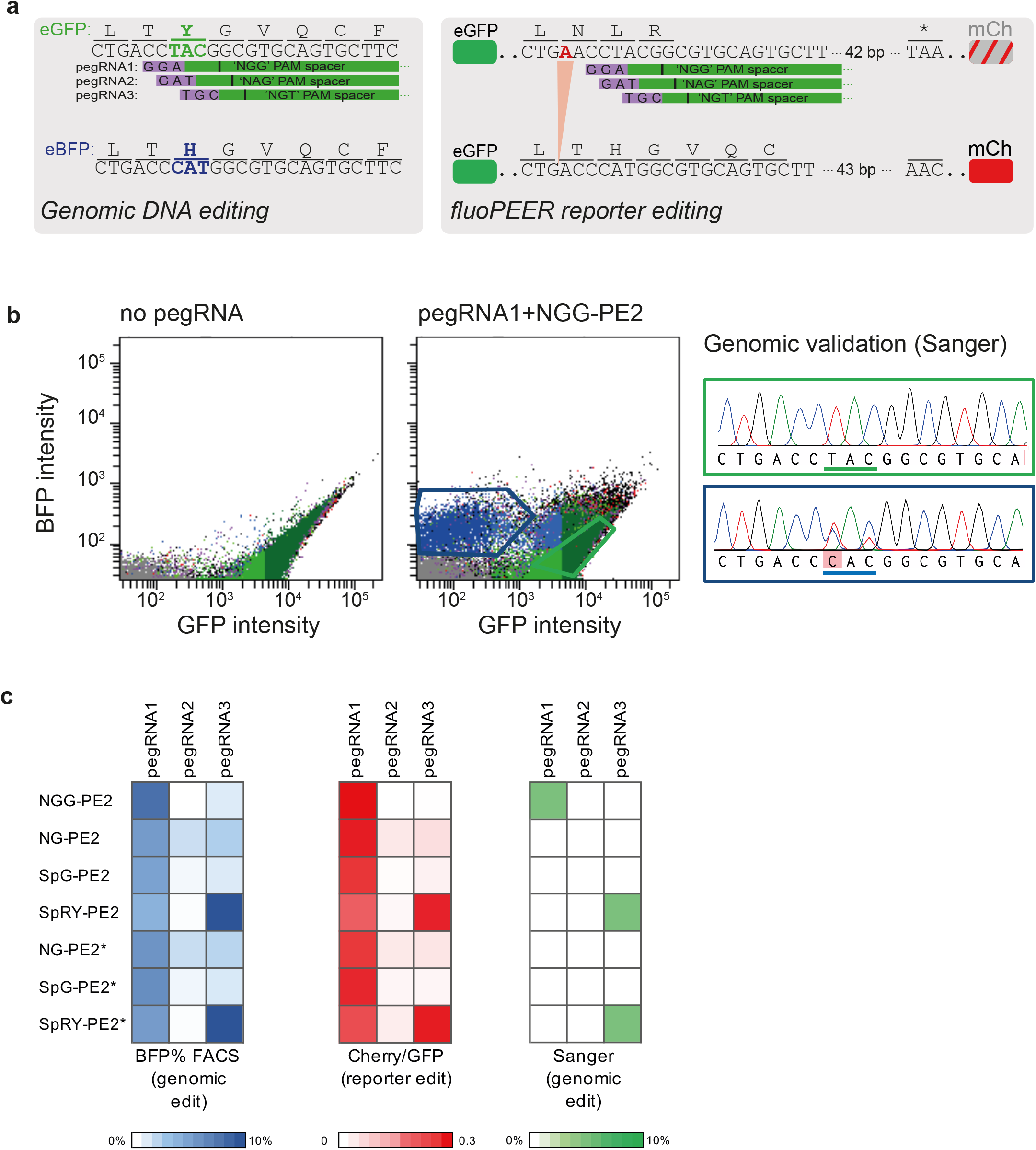
Comparison of genomic editing of GFP→BFP to editing on the corresponding fluoPEER plasmid. **a**, Design of three different pegRNAs converting GFP to BFP (TAC to CAT). The same three pegRNAs remove a single nucleotide insert on the corresponding fluoPEER plasmid, resolving a frameshift mutation and upstream of the Cherry cassette. Note that the PAM sequences (purple) of the pegRNA 1, 2, and 3 are NGG, NGN, and NAN, respectively. **b**, HEK293T cells containing a lentivirally integrated genomic GFP cassette were transfected with the prime editing machinery to convert GFP to BFP. FACS plot shows occurrence of BFP^+^ cells 14 days after transfection with pegRNA1 (NGG-PAM) and NGG-PE2. Sanger sequencing confirms TAC to CAT conversion in BFP^+^ cells (blue outlines in FACS and Sanger), but not in GFP^+^ cells (green outlines). **c**, Editing of integrated genomic GFP (left panel) and editing of the corresponding fluoPEER plasmid (middle panel) in HEK293T cells using the three pegRNA designs from (a) and 7 different PE2 variants (‘no PE2’ is included in Fig 1d as an additional PE2-condition). Sanger sequencing of the genomic target region (right panel) in the same cells as shown in the left panel illustrates that this technique is not sensitive enough to quantify GFP to BFP conversion in less than 5% of cells. Related to figure 1d.

**Supplementary Figure 5:**
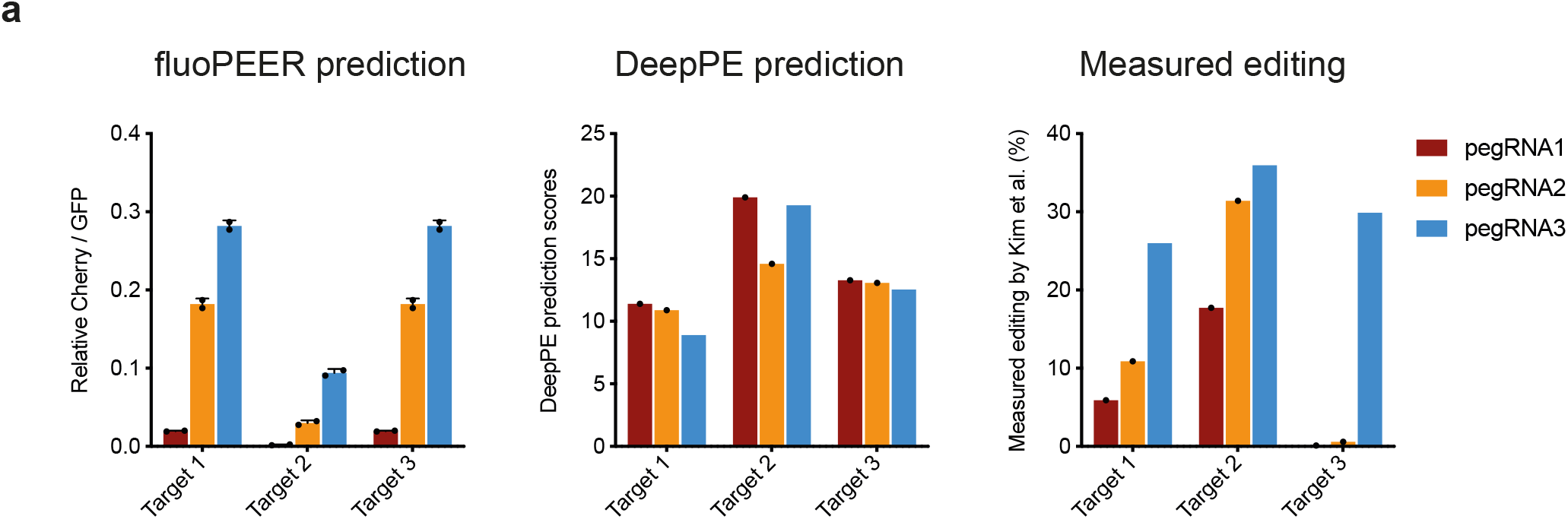
Comparison of fluoPEER ranking to predictions by the DeepPE algorithm. **a**, Comparison of the ranking of editing efficiencies by fluoPEER (left panel) to the prediction score of the DeepPE algorithm (middle panel) and to actual prime editing efficiencies as reported by Kim et al., 2021. (right panel).

**Supplementary Figure 6:**
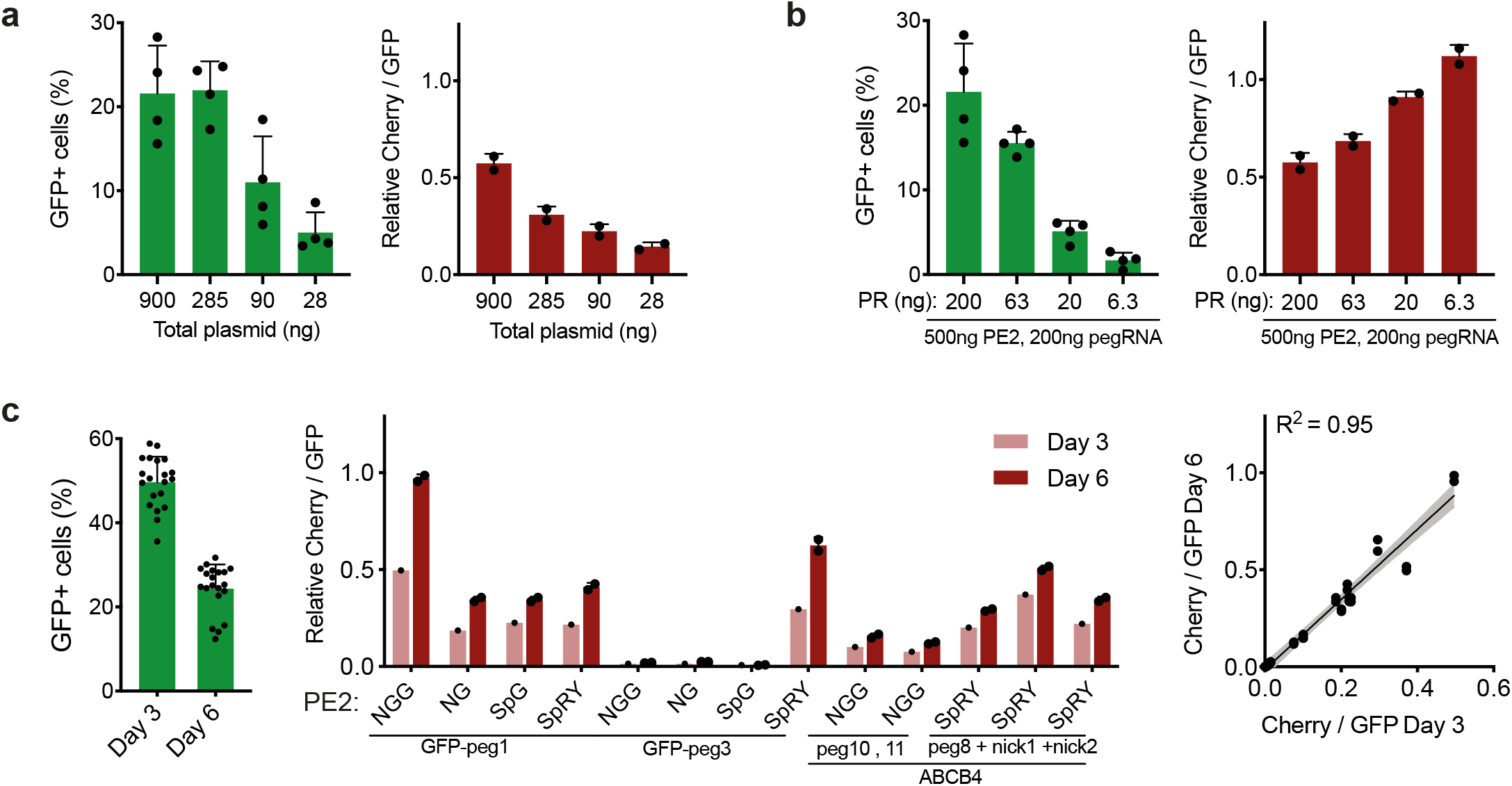
Dynamics of the fluoPEER under various transfection conditions and read-out times. **a**, Percentage of GFP+ cells and average of Cherry to GFP ratio in GFP+ HEK293T cells, with decreasing amounts of total transfected plasmid DNA (PE2+pegRNA+PR). **b**, Percentage of GFP+ cells and average of Cherry to GFP ratio in GFP+ HEK293T cells, with decreasing amounts of PR plasmid DNA but equal amounts of PE2 and pegRNA plasmid DNA. As expected, lowering PR led to decreased percentages of GFP+ cells, but increased reporter editing (Cherry to GFP ratio). **c**, Percentage of GFP+ cells and average of Cherry to GFP ratio in transfected cells, as measured 3 and 6 days after transfection of PE2+pegRNA+PR. Note that reporter editing increases linearly over time. Therefore, measuring Cherry to GFP ratio at day 3

**Supplementary Figure 7:**
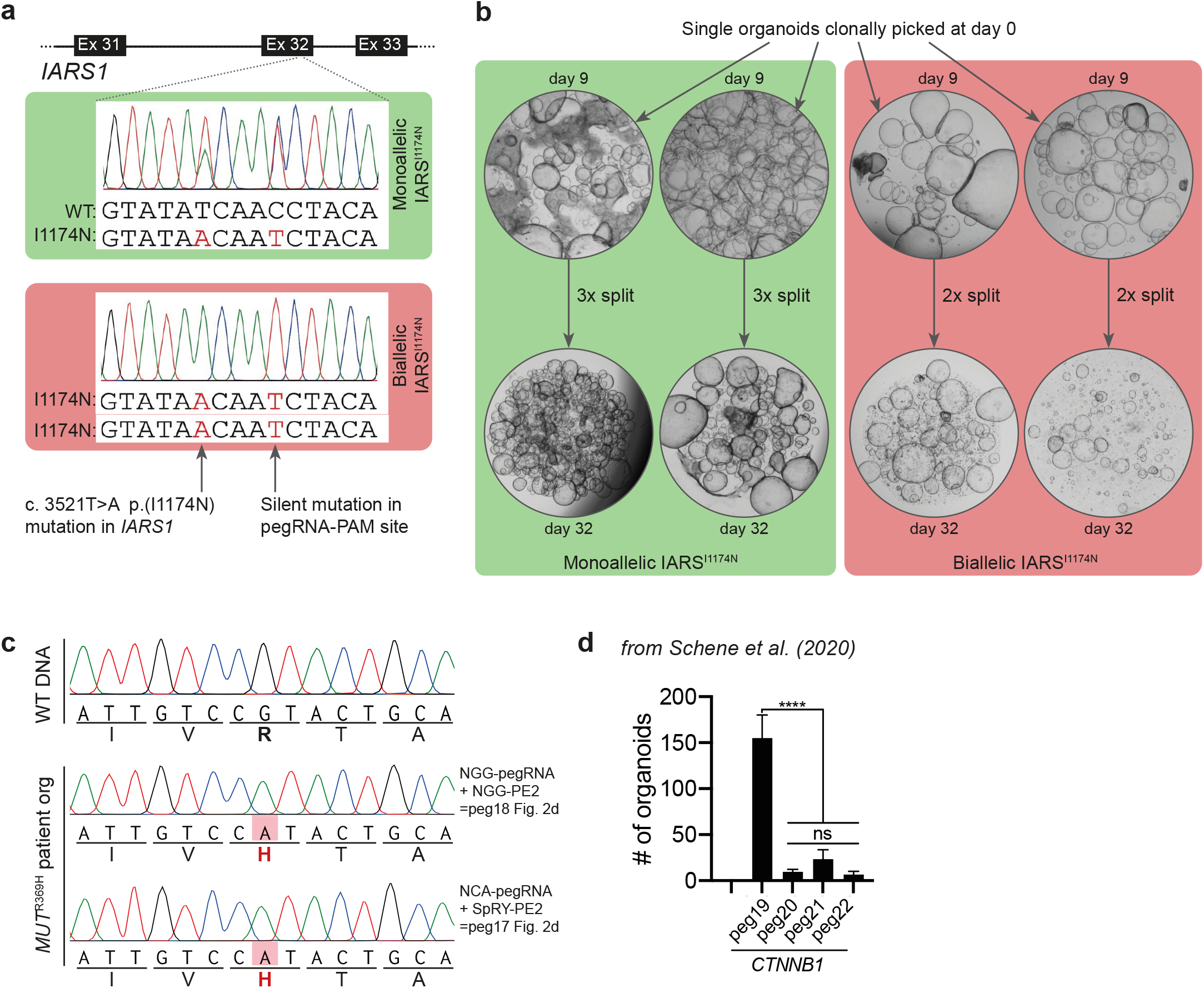
fluoPEER-predicted prime editing in organoids. **a**, Sanger sequencing of clonal liver-derived organoids with monoallelic and biallelic I1174N mutations in exon 32 of *IARS1*. **b**, Clonal organoid lines with monoallelic (2 clones) and biallelic (2 clones) *IARS1*^I1174N^ mutations were continually passaged for 32 days. Clones with biallelic, but not monoallelic, IARS^I1174N^ mutations had lower organoid reconstitution capacity and could be passaged less often due to slower growth. **c**, Correction of pathogenic *MUT*^R369H^ mutations in liver organoids derived from an MMA patient could not be corrected using two different pegRNA designs. This corresponds to very low fluoPEER prediction scores for these targeting strategies as shown in Fig. 2e. **d**, Organoid reconstitution in medium without Wnt-pathway agonists Rspo1 or Wnt, after prime-editing mediated introduction of stabilizing *CTNNB1* mutations that cause Wnt-independent growth. Note that pegRNA19 significantly outperformed other pegRNA designs in both the fluoPEER analysis (Fig. 2d) and genomic editing of *CTNNB1*. Data in (d) was adapted from earlier experiments within our lab (Schene et al. 2020).

**Supplementary Figure 8:**
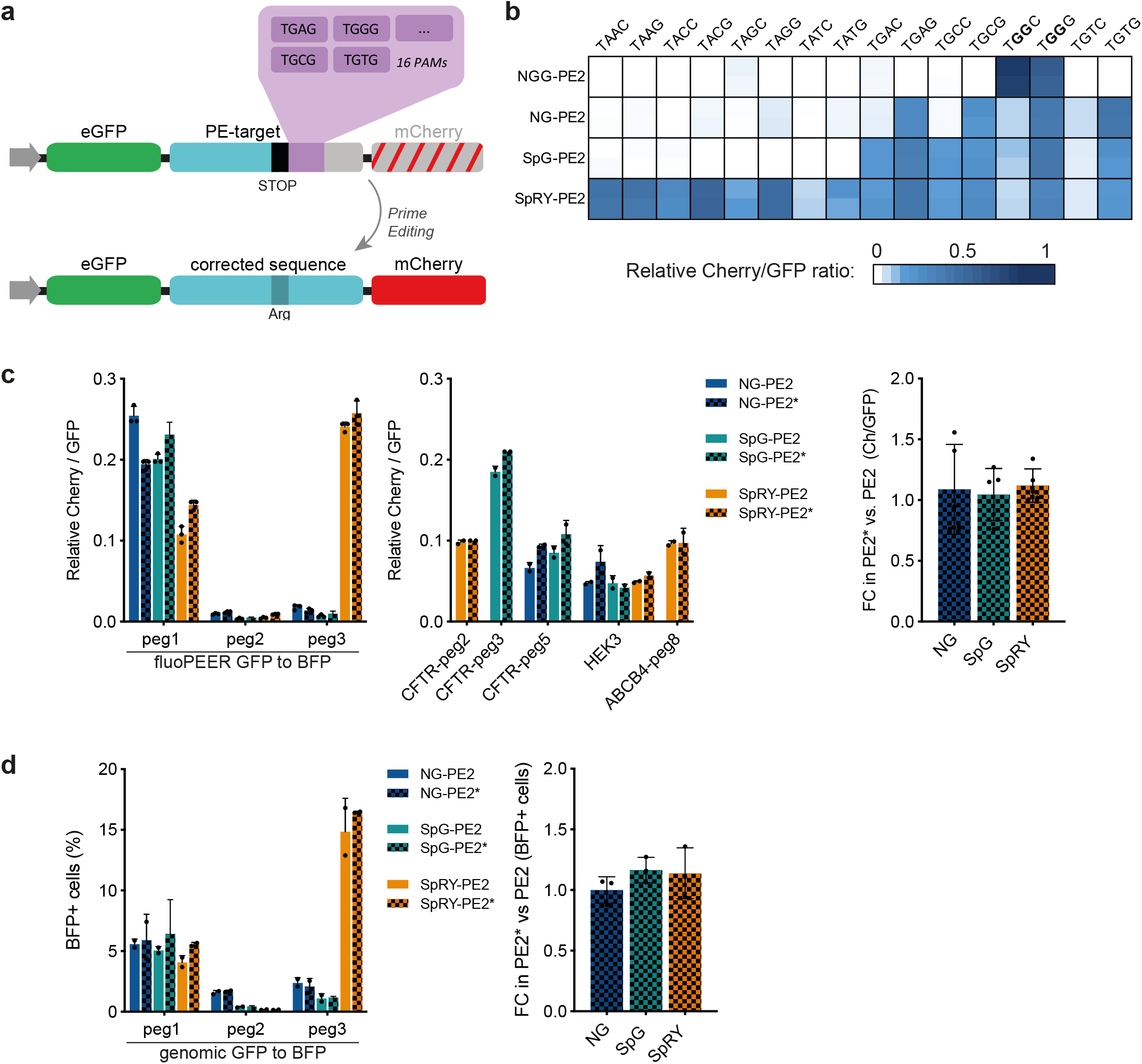
Characterization of PAM-specificity and NLS adaptations of novel PE2 variants using fluoPEER. **a**, Schematic of the design of 16 different fluoPEER plasmids, each containing the same pegRNA spacer binding site (HEK3) containing a stop codon at position 18-20 followed by a variable 4-nucleotide PAM sequence. Prime editing using a pegRNA that converts this stop codon to an arginine-encoding codon results in Cherry signal. **b**, Heatmaps of Cherry to GFP ratio for the 16 plasmids in (a), using four PE2 variants. We find that PAM-specificities of PE2 variants are highly similar to those described for corresponding cutting Cas9 variants (Walton et al., 2020). Note that a guanine (G) nucleotide on position 4 of the PAM sequence is associated with higher overall editing efficiency. **c**, The left panels show a comparison of fluoPEER scores for flexible PE2 variants and the corresponding PE2* variants with nuclear localization sequence (NLS) adaptations as described by Liu et al. (2021). The right panel shows a summary of all fluoPEER scores, expressed as the fold change (FC) of the PE2* variants relative to the corresponding PE2 variants. **d**, The left panel shows conversion of genomic GFP to BFP using various PE2 and PE2* variants. The right panel shows a summary of the left panel, expressed as the FC between the PE2* and the PE2 variant. Note that the PE2* NLS adaptations did not result in an overall increase of fluoPEER scores nor an overall increase in genomic editing efficiency. The data shown in Supplementary Fig. 4c was reused for (c) and (d).

**Supplementary Figure 9:**
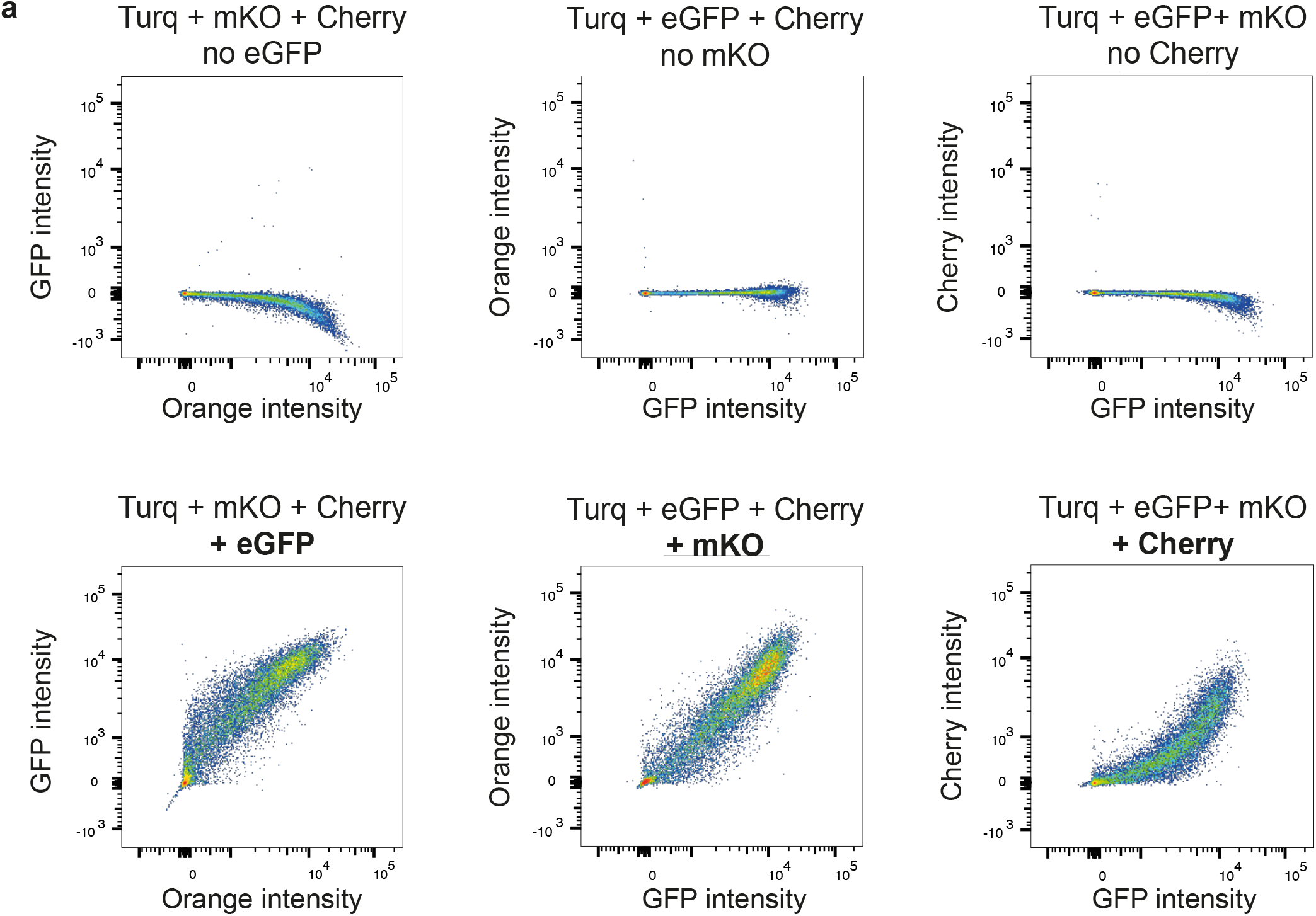
Dynamics of cotransfection of four fluorescent plasmids. **a**, Mixtures of plasmids (5kb) containing CMV-mTurquoise2, CMV-eGFP, CMV-mKO2, and/or CMV-mCherry were transfected into HEK293T cells and inspected by FACS after 2 days. FACS plots show proper compensation in the ‘fluorescence minus one’ (FOI) controls and strong correlation between fluorescent intensities from different plasmids in the condition containing all 4 plasmids, indicating very high cotransfection efficiency. This strongly suggests that 2-4 fold enrichment of genomic editing in GFP^+^Cherry^+^ versus GFP^+^Cherry^-^ cells (Fig. 2a) cannot be the result of inefficient cotransfection of plasmids encoding prime editing machinery (PE2+pegRNA) in GFP^+^Cherry^-^ cells.

**Supplementary Figure 10:**
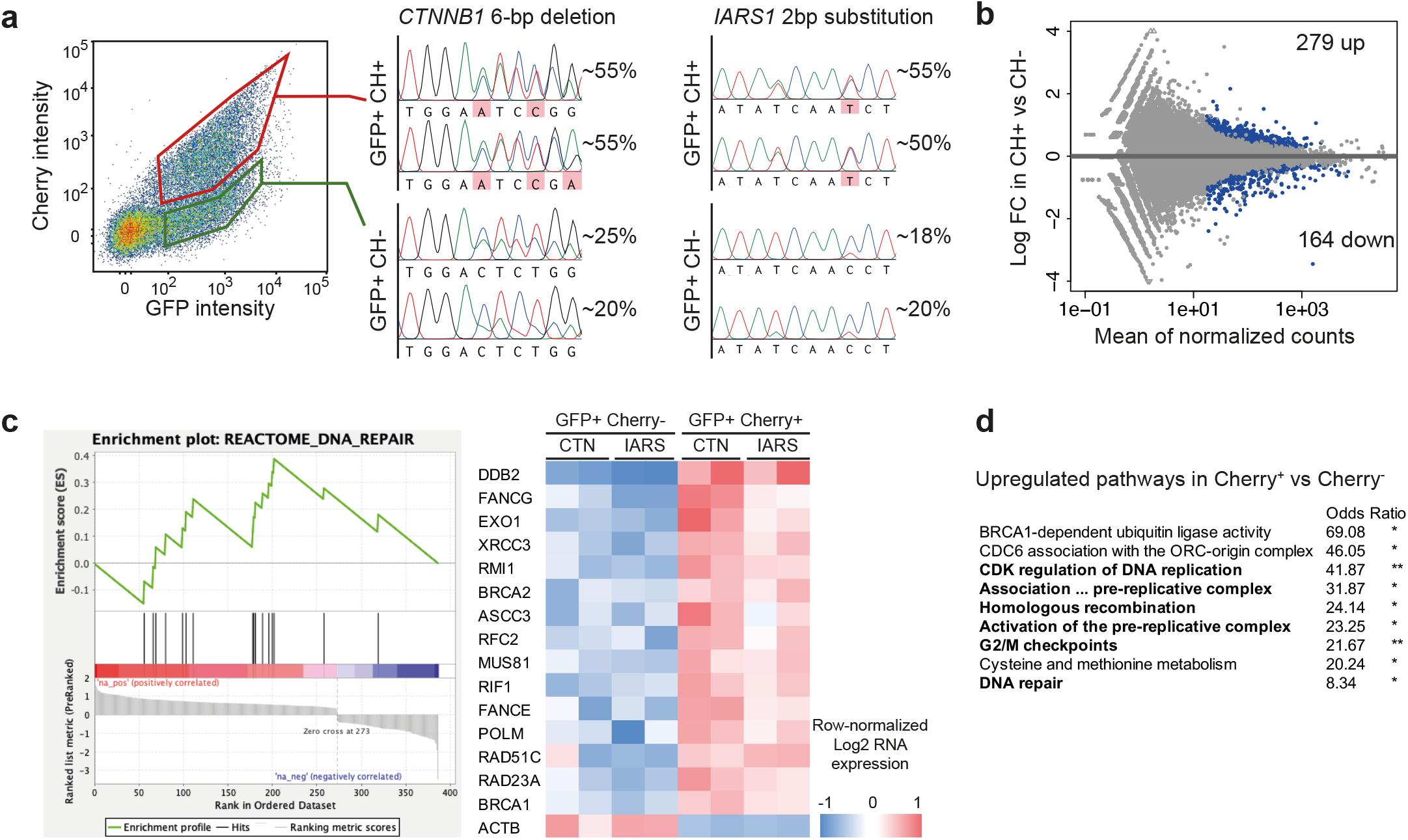
RNA sequencing analysis of fluoPEER-edited versus - unedited cells. **a**, HEK293T cells were transfected with PE2 and pegRNA plasmids creating either IARS^I1174N^ mutations or *CTNNB1* 6-bp deletions, together with corresponding fluoPEER plasmid. 2 days after transfection, GFP^+^Cherry^+^ and GFP^+^Cherry^-^ cells were sorted and genomic editing was estimated by Sanger sequencing. **b**, RNA-sequencing was performed in replicates of each population from (a). MA-plot shows significant (adjusted *p*-value <0.05) up- and downregulation of 279 and 164 genes, respectively in GFP^+^Cherry^+^ versus GFP^+^Cherry^-^ cells. **c**, The left panel shows the enrichment plot of the most enriched ‘Reactome Database’ pathway in GFP^+^Cherry^+^ versus GFP^+^Cherry^-^ cells. The right panel shows the expression heatmap of enriched genes within this pathway. **d**, EnrichR analysis of GFP^+^Cherry^+^ versus GFP^+^Cherry^-^ cells.

## Methods

### Study approval and human subjects

The study was approved by the responsible local ethics committees (Institutional Review Board of the University Medical Center Utrecht and University Medical Center Groningen (STEM: 10-402/K; TcBio 14-008; Metabolic Biobank: 19-489)). For cystic fibrosis organoids, collection of patient tissue and data was performed following the guidelines of the European Network of Research Ethics Committees (EUREC). Tissue biopsies from the liver of a patient with ABCB4 deficiency (PFIC3) was obtained during a liver transplant procedure in the UMCG, Groningen. Rectal biopsies used for intestinal organoid culture from a patient with ATP8B1 deficiency (PFIC1) were obtained at the outpatient clinic in the UMCU, Utrecht. Biobanked intestinal organoids are stored and cataloged (https://huborganoids.nl/) at the foundation Hubrecht Organoid Technology (http://hub4organoids.eu). All biopsies were used after written informed consent.

### Plasmid cloning

fluoPEER plasmids were cloned using the backbone of the pmGFP-P2A-K0-P2A-RFP (Addgene #105686) plasmid, which was a gift from Ramanujan Hegde. This plasmid was cut with SalI and Acc65I for 16 hours at 37 degrees, creating ‘TCGA’ and ‘GTAC’ overhangs. Genomic insert oligos containing 5’ ‘TCGACC’ and 3’ ‘G’ overhangs on the top oligo, and 5’ ‘GTACC’ and 3’ ‘GG’ overhangs on the bottom oligo were annealed and inserted using a cycling protocol (see supplementary information 1).

Cloning of pegRNA plasmids was performed according to previously described protocols (Anzalone, 2019). In brief, the pU6-pegRNA-GG-Vector (Addgene #132777) was digested for 16 hours with BsaI-HFv2 (NEB), after which the 2.2 kb fragment was isolated from gel. Oligonucleotide duplexes of the pegRNA spacer, pegRNA extension, and pegRNA scaffold sequences were ordered containing the appropriate overhangs and subsequently annealed. The annealed pegRNA duplexes were ligated into the pU6-pegRNA-GG-Vector using Golden Gate assembly with BsaI-HFv2 (NEB) and T4 DNA ligase (NEB) in a protocol of 12 cycles of 5 min at 16 °C and 5 min at 37 °C. For cloning of sgRNAs used for PE3 and ABE-NG, we replaced the BsmBI restriction sites of the BPK1520 plasmid by BbsI restriction sites using PCR, which allowed direct ligation of sgRNA-spacer duplexes (Ran, 2013). All pegRNA, sgRNA, and primers sequences used in this work are listed in Supplementary Table 1 and were synthesized by Integrated DNA Technologies (IDT). pCMV-PE2 (Addgene #132775) and pU6-pegRNA-GG-acceptor (Addgene #132777) were gifts from David Liu; BPK1520 (Addgene #65777) was a gift from Keith Joung.

### Cloning of flexible PE2s and PE2*

Using PCR and In-Fusion cloning (Takara Bio), the ‘NGG’ PAM-recognition domain of the prime editor protein (PE2) was replaced with the corresponding domains in NG-ABEmax (Huang et al., 2019), SpG-ABEmax, or SpRY-ABEmax (Walton et al., 2020). NG-ABEmax was a gift from David Liu (Addgene #124163). SpG- and SpRY-ABEmax were a gifts from Benjamin Kleinstiverwas (Addgene plasmids #140002 and #140003). PE2* variants with improved nuclear localization sequence (NLS) were adapted from the NGG-PE2* developed by Liu et al. (2021) and were cloned by PCR and In-Fusion cloning (Takara Bio).

### Organoid culture

Liver and intestinal organoids were cultured according to previously described protocols. In short, liver organoids were plated in matrigel (Corning) and maintained in liver expansion medium (EM), consisting of AdDMEM/F12 (GIBCO) supplemented with 2% B27 without vitamin A (GIBCO), 1.25 mM N-Acetylcysteine (Sigma), 10 mM Nicotinamide (Sigma), 10 nM gastrin (Sigma), 10% RSPO1 conditioned media (homemade), 50 ng/ml EGF (Peprotech), 100 ng/ml FGF10 (Peprotech), 25 ng/ml HGF (Peprotech), 5 mM A83-01 (Tocris), and 10 mM FSK (Tocris). Small intestine and colon organoids were plated in matrigel and maintained in SI-EM, consisting of AdDMEM/F12 supplemented with 50% WNT3A-, 20% RSPO1-, and 10% NOG(gin)-conditioned medium (all homemade), 2% B27 with vitamin A (GIBCO), 1.25 mM N-Acetylcysteine, 10 mM Nicotinamide, 50 ng/ml murine-EGF (Peprotech), 500 nM A83-01, and 10 mM SB202190 (Sigma). The medium was changed every 2–4 days and organoids were passaged 1:4 –1:8 each week. After thawing, organoids were passaged at least once before electroporation.

### Cell culture

HEK293T cells were maintained and split every 4-5 days. For production of lentivirus, HEK293T cells were plated in a 145mm CELLSTAR dish (Corning) and transfected 24 hours later (at 50-60% confluence) with a mix of 10 μg of the pLenti-CMV-GFP-Hygro plasmid, 5 μg of psPAX2, 5 μg of pMD2.G, and 60 μl of polyethylenimine (1 mg/ml). pLenti-CMV-GFP-Hygro was a gift from Eric Campeau & Paul Kaufman (Addgene #17446), psPAX2 and pMD2.G were gifts from Didier Trono (Addgene plasmid # 12260 and # 12259). 24 hours after transfection, the medium was replaced with DMEM containing pen/strep supplemented with 10% (vol/vol) FBS, and virus containing medium was harvested at 48 and 96 hours after transfection. Medium was centrifuged at 1,000 r.p.m. for 5 min and supernatants were filtered through a 0.22-μm filter, after which virus particles were concentrated by ultracentrifugation at 50,000 x g for 2 hours and resuspension in 1 ml DMEM. HEK293T cells were transduced at an MOI of 2 for 24 hours in the presence of polybrene (8 μg/μl) and analyzed by FACS 14 days after transduction to confirm stable GFP expression. For transfection, HEK293T were split at 50.000 cells per well in a 96 wells plate one day prior to transfection. HEK293T cells were transfected with 0.1 μg fluoPEER, 0.25 μg PE2 plasmid, 0.1 μg pegRNA plasmid and 0.05 μg nicking-gRNA plasmid in a mix of 25 μl OptiMEM and 0.5 μl lipofectamine 2000 for each well.

### Electroporation

Before electroporation, organoids were grown under standard culture conditions. Four wells containing organoids were then dissociated for each condition using TrypLE for 4–5 min at 37 °C, after which mechanical disruption was applied through pipetting. Cells were washed once using Advanced DMEM/F12, resuspended in 80 μl OptiMEM containing Y-27632 (10 μM), and 20 μl DNA mixture was added. For prime editing, the DNA mixture contained 5 μg fluoPEER, 12 μg PE2 plasmid, 5 μg pegRNA plasmid and 2 μg nicking sgRNA plasmid. The cell-DNA mixture was transferred to an electroporation cuvette and electroporated using a NEPA21 electroporator (NEPA GENE) with 2× poring pulse (voltage: 175 V, length: 5 ms, interval: 50 ms, polarity: +) and 5× transfer pulse (voltage: 20 V, length: 50 ms, interval: 50 ms, polarity ±), as previously described (Fujii, 2015). Cells were removed from the cuvette and transferred into 500 μl OptiMEM containing Y-27632 (10 μM). After 20 minutes, cells were plated in 180 μl matrigel divided over 6 wells. Upon polymerization of the Matrigel, medium was added containing Y-27632 (10 μM).

### FACS

Organoids and cell lines were harvested and digested to single cells, after which cells were resuspended in FACS buffer (phosphate-buffered saline with 2 mM ethylenediaminetetraacetic acid and 0.5% bovine serum albumin). Prior to FACS, cells were filtered through a 5 ml Falcon polystyrene test tube (Corning). Flow cytometry was performed on the FACS Fortessa (BD) and sorting was performed on the FACS FUSION (BD) using FACS Diva software (BD). Sorted cells were collected in culture medium and spun down.

### Genotyping

Sorted cells were harvested using the Quick-DNA microprep kit (Zymogen) according to manufacturer’s protocols. PCR was performed on the genomic region of interest using the Phusion polymerase (ThermoFisher) and purified using the <u>QIAquick PCR Purification Kit</u> (Qiagen) according to manufacturer instructions. The PCR product was sent for Sanger sequencing to EZSeq Macrogen Europe.

### RNA sequencing

mRNA was isolated using Poly(A) Beads (NEXTflex). Sequencing libraries were prepared using the Rapid Directional RNA-Seq Kit (NEXTflex) and sequenced on a NextSeq500 (Illumina) to produce 75 base long reads (Utrecht DNA Sequencing Facility). Sequencing reads were mapped against the reference genome (hg19 assembly, NCBI37) using BWA (Li and Durbin, 2009) package (mem –t 7 –c 100 –M –R). Raw reads were further analyzed as described under ‘Data analysis’.

### Cell cycle analysis

200 ng/ml nocodazole was added to HEK293T cells 24 hours before FACS analysis as a cell cycle control. 24 hours before fluoPEER read-out and genomic sanger sequencing, HEK293T cells were transfected. 48 hours before RNA isolation, HEK293T cells were transfected.

### Data analysis

Flow cytometry data was analyzed using FlowJo. RNA sequencing was analyzed using DESeq2 in RStudio (Love, 2014), gene set enrichment analysis (Subramanian, 2005), and enrichR (Chen, 2013). All figures were made in Prism GraphPad or GGPlot2 (Wickham, 2012) in RStudio. Sanger sequencing was quantified using EditR (Kleusner, 2018) or TideR (Brinkman, 2018).

### ClinVar database computational analysis

Information for all pathogenic mutations shorter than 51 bp was obtained from the ClinVar database (Landrum, 2018), accessed October 2020. Genomic sequences flanking these mutations were obtained from RefSeq (Pruitt, 2005) accessed October 2020, using the SPDI data model (Holmes, 2020) and a custom python script. The −10 to +4 bp region around the target sites were searched for NGG, NAN or NGN PAMs. The efficiency of prime editing using NGG PAMs was predicted using PE_Position and PE_type random forest models, provided by (Kim, 2021). Figures were made in python using Matplotlib (Hunter, 2007). The code used for this analysis is available on GitHub (https://github.com/JBaijens/PE_prediction).

## Supplementary Note 1: fluoPEER cloning

### STEP 1

Digest the pmGFP-P2A-K0-P2A-RFP plasmid by making the following mix:

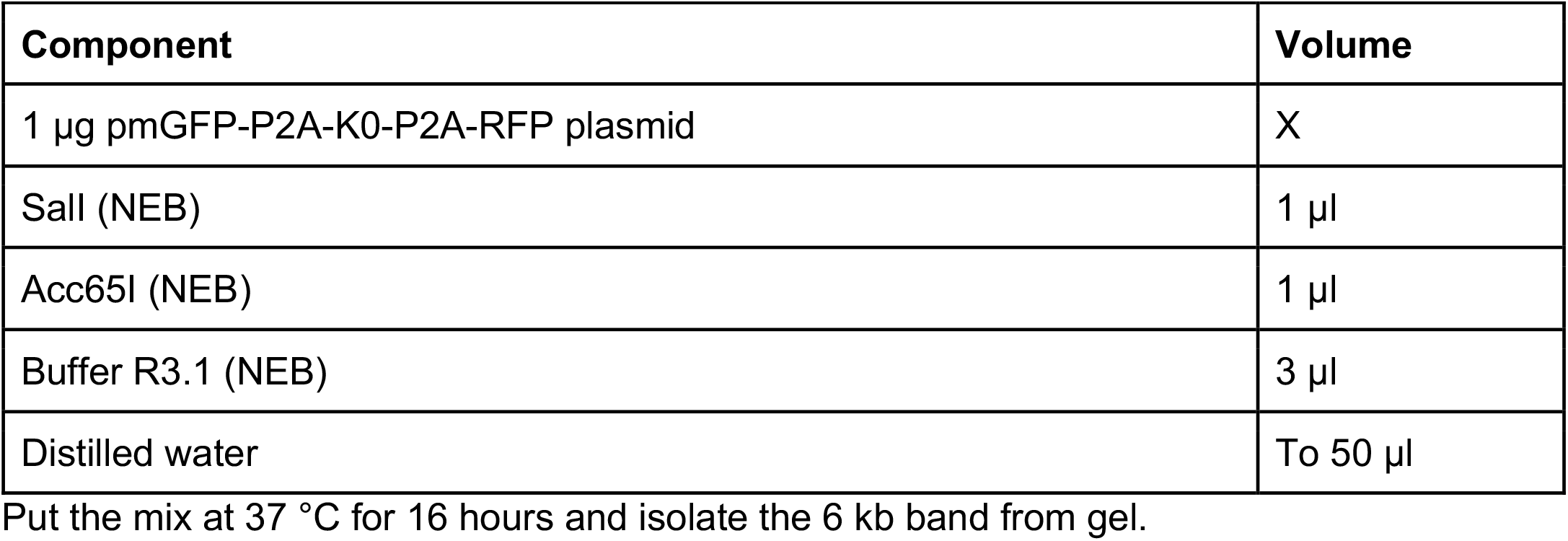

fluoPEER needs a STOP codon or frameshift in the genomic insert to function. In case the genomic target already contains either, 45-100 bp around the mutation of interest can be selected. The genomic insert should at least contain the full spacer and extension (PBS+RTT) sequences of all designed pegRNAs, including a margin of 5bp upstream and 5bp downstream. In case none such frameshift nor STOP-codon is present, one will have to be inserted. Important notes:

- The fluoPEER plasmid should be the one modified, such that the pegRNA can be used on both the fluoPEER plasmid and the genomic target.
- The PAM site and spacer sequence (including PBS) should not be altered by addition of frameshift or creation of stop-mutation. In case this is unavoidable, design two fluoPEER plasmids for the mutation instead.

### STEP 2

Once the genomic insert has been selected, two oligos need to be ordered: TOP: 5’TCGACC’-[GENOMIC INSERT]-’G’3 BOTTOM: 5’GTACC’[GENOMIC INSERT REVERSE COMPLEMENT]-’GG’3

Anneal the oligos according to the following protocol:

| Component | Volume |
| --- | --- |
| TOP oligo (100 µM) | 1 µl |
| BOTTOM oligo (100 µM) | 1 µl |
| PNK (NEB) | 1 µl |
| PNK buffer 10x (NEB) | 1 µl |
| Distilled water | 6 µl |
| **Total** | 10 µl |

Heat the mix up to 95 °C for 3 minutes and ramp down to 20 °C at 5 °C / min.

Dilute the oligos 1:10 by adding 90 μl water to the mix.

### STEP 3

Ligate the plasmid and annealed oligos in the following mix:

| Component | Volume |
| --- | --- |
| Digested pmGFP-P2A-K0-P2A-RFP plasmid | 30 ng |
| Annealed genomic insert | 1 μl @ 1 μM |
| T4 DNA ligase (NEB) | 1 μl |
| T4 DNA ligase buffer 10X (NEB) | 1 μl |
| Distilled water | To 10 μl |

Set at 21 °C for 60 minutes and transform.

